# Comparison of microbial and sono-assisted alkaline pre-treatment of sugarcane bagasse, followed by one-pot synthesis of microbial extracellular polymeric substance through simultaneous delignification, saccharification, and fermentation

**DOI:** 10.1101/2024.06.22.600045

**Authors:** Ankita Debnath, Sundipan Bhowmick, Ramkrishna Sen

## Abstract

Lignocellulosic biomass (LCB) captures a major fraction of agro-industrial wastes that are mostly valorised for the production of second-generation biofuels. The extensive pre-treatment followed by saccharification of LCB restricts its usability for the production of a wide array of bioproducts. This study highlights the performance comparison of sono-assisted alkaline pre-treatment versus microbial pre-treatment of sugarcane bagasse by a no. of analytical techniques such as FTIR, XRD, CHNS, and SEM. Moreover, simultaneous delignification, saccharification and fermentation (SDSF) in one pot is highly desirable for the cost effective and environment friendly production of microbial products. In the present study SDSF was carried out by a cellulolytic bacterium *Cellulomonas flavigena,* for the production of extracellular polymeric substance (EPS), which is, by chemical nature a biopolymer made up of carbohydrate and protein subunits. The biopolymer was found to have an overall positive charge as analysed by ion exchange chromatography. Two major fractions of molecular weight 237 KDa and 29 KDa were obtained from gel filtration chromatography, in addition to that, the EPS was found to be composed of monosaccharides, D (+) mannosamine and D (+) xylose as identified from high pressure liquid chromatography (HPLC).

## Introduction

Sugarcane bagasse have been mostly utilized in biotechnology for the production of biofuels (Tsegaye et al., 2019; Velmurugan and Muthukumar, 2012). Bioethanol industry have mostly been focussing on 2G biofuels, which are produced from lignocellulosic biomass (LCB) such as rice husk, wheat straw, sugarcane bagasse, wood, rice straw among others (Velmurugan and Muthukumar, 2012). Although 2G biofuels are produced from these agro-industrial wastes, which are quite abundantly available in countries like India, still this idea suffers due to the presence of lignin in these wastes. Lignin is a major component of LCB, that is responsible for the recalcitrant nature of cellulose and hemicellulose in LCB. The enzymatic saccharification of cellulose becomes low due to limited accessibility of these polysaccharides as a result of the lignin masking them. Thus, production of biofuels from LCB requires extensive pre-treatment of the biomass to remove lignin so that the polysaccharides become available for hydrolysis. The hydrolysate obtained after saccharification is then used for fermentation by yeasts to produce biofuels. The lignin removal and further saccharification of LCB by alkaline or acid pre-treatment and hydrolysis not only increases the overall cost of operation but also leaves behind furfural like derivatives that are detrimental for microbial viability during fermentation. To avoid the extra cost and formation of harmful derivatives, biological pre-treatment methods like fungal cellulase and hemi-cellulase mediated saccharification preceding the lignin removal can be widely found in literature (Maeda et al., 2011). For example, cellulase from *Trichoderma* have been used in various studies for the saccharification of cellulose after removal of lignin by acid or alkaline pre-treatment (Delabona et al., 2012; Maeda et al., 2011) . Moreover, the usage of LCB has been confined mostly for the production of biofuels. Reports of fermentation of other value-added microbial products like bioplastics, biopolymers like bacterial extracellular polymeric substances (EPS) are rarely found in literature (Andharia et al., 2023; Terán Hilares et al., 2019). EPS or bacterial extracellular polymeric substances are mixture of proteins and polysaccharides, produced in huge quantities inside the bacterial cells, which are then transported outside by the cells as combination of extracellular macromolecules (Rabiya and Sen, 2022). EPSs like pullulan, curdlan, hyaluron etc. have been proved to be of immense industrial importance from environmental to biomedical sectors (Gupta et al., 2024), thereby requiring cost-effective production of these biopolymers from waste resources. Although some reports of PHA production from LCB can be found but EPS production by fermenting LCB is hard to find. This is most likely because of the cost involved in delignification by chemical or chemical plus enzymatic pre-reatments in tandem (Thite and Nerurkar, 2019). To avoid extra steps of pre-treatment followed by several washings (Bhowmick., 2023), this study directly used raw sugarcane bagasse for saccharification by a cellulolytic bacterium *Cellulomonas flavigena* (CF), which subsequently fermented the resulting reducing sugars from hydrolysed SCB to produce EPS.

In the present study agro-industrial waste product sugarcane bagasse was used as a feedstock for the production of extracellular polymeric substance (EPS) by a common cellulolytic bacterium *Cellulomonas flavigena* using the concept of consolidated bioprocessing. *Cellulomonas flavigena* have been much studied for its cellulolytic potential (Maki et al., 2009) but studies involving single step one pot fermentation of LCB using this bacterium are few. This idea of consolidated bioprocessing was incorporated in the present study to have an environment friendly alternative towards future environmental engineering applications for example in self-healing of concrete and soil stabilization, with the idea of using this EPS as an encapsulating or coating material for biomineralizing bacteria. In addition to having a cleaner production of bacterial EPS, this study also comprises of a comparative analysis of sono-assisted alkaline and microbial pre-treatment by *Cellulomonas flavigena*. Thus, this study is a unique approach towards performance assessment of physico-chemical vs biological pre-treatment and consolidated one pot saccharification and fermentation of sugarcane bagasse to produce bacterial EPS along with the partial characterization of the bacterial extracellular product.

## Materials and methods

### Performance evaluation of microbial depolymerization of sugarcane bagasse (SCB) in terms of delignification and saccharification

Sugarcane bagasse was collected from local market at IIT Kharagpur washed, dried and grinded. The ground bagasse was sieved in ASTM test sieves of size 1.15mm. The bagasse particles were then directly used for bacterial delignification followed by saccharification and fermentation. Sugarcane bagasse was used as the sole carbon source in M9 minimal media and were subjected to simultaneous saccharification and fermentation by cellulolytic bacterium *Cellulomonas flavigena* NCIM Accession no. 5604 obtained from NCIM Pune, India. Culture was maintained at nutrient agar medium pH 7, 30°C. For microbial pre-treatment 1% (w/v) of sugarcane bagasse as the sole carbon source in M9 minimal media composed of Na_2_HPO_4_.2H_2_O 4.2 g/l, KH_2_PO_4_ 1.5 g/l, NaCl 0.25 g/l, NH_4_Cl 0.5 g/l yeast extract 0.5 g/l. The bacterial depolymerization was carried out till 120 hours, followed by the characterization of the digested biomass, in terms of elemental analysis, functional groups, crystallinity and morphology.

### Performance comparison between microbial depolymerization and sono-assisted alkali treated deconstruction of SCB

For sono-assisted alkaline pre-treatment of sugarcane bagasse (SCB) various ratios of SCB: alkali was maintained with a specific heating temperature. The ratios maintained were at 1:20, 1:25, 1:30 and 1:35 LSR (liquid solid ratio). Pre-treatment conditions were 2% w/v NaOH with 1% w/v sodium chlorite at 95°C heating for 1 hour followed by 4 hours of heating at 50°C, with continuous stirring. The chemical pre-treatment was followed by 1.5 hours of sonication with 45 secs pulse on and 15 secs pulse off at 50 hz frequency. Before analysis all the samples were neutralized by continuous washing with distilled water. After the physio-chemical pre-treatment samples were lyophilized for performance evaluation in terms of delignification and depolymerisation.

### Characterization of microbially and chemically treated SCB Physical characterization

#### FTIR

The freeze-dried bagasse particles subjected to FTIR analysis after mixing with KBr pellets, the spectra were recorded from 400 to 4000 cm^-1^ in a Nicolet 670 Fourier Transform Infrared Spectrometer, spectrum was recorded out of 32 scans for each sample

#### XRD

A Bruker D8 Advance X-ray diffractometer (XRD) with Eiger2 R 500K detector 0/90° with Cu Kα (λ = 1.54 Å) radiation source was used for bagasse particles, the scan range was kept at 10-50°. The crystallinity index of all the samples were quantified by the following formula (Bhowmick et al., 2023):

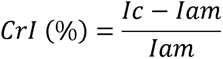

Here, I_c_ represents peaks of crystalline cellulose at plane 002 at 20 of 22.5°C, I_am_ represents amorphous peaks of cellulose between 200 and 110 plane of 20 at 18°C.

#### Elemental composition by CHNS

C, H, N, S and inorganic ash content was analysed using a CHNS elemental analyser. This elemental analysis was essential to evaluate the delignification and de-polymerization of sugarcane bagasse by chemical and microbial treatment.

### Microscopic characterization by SEM

The morphological characterizations were performed by SEM (ZEISS) equipped with a GEMINI II column. Before observing under SEM samples were dried overnight in a vacuum desiccator and gold coated.

### Microbial growth on pretreated and untreated sugarcane bagasse

Growth studies of *Cellulomonas flavigena* on pretreated and 1% (w/v) untreated sugarcane bagasse were performed using Triphenyl tetrazolium chloride (TTC) reagent, which results in the formation of pink coloured product formazan if metabolically active cells are present in a culture (Defez et al., 2017). The media components, M9 minimal medium, composition mentioned in previous section. Absorbance of formazan measured at 480 nm was taken as a direct measure of growth. The total carbohydrate produced by the cells as result of delignification, saccharification were also monitored by phenol sulphuric acid assay by maintaining the ratio of culture supernatant: phenol: sulphuric acid as 1:1:5 (Dubois et al., 1956). For getting the concentration of carbohydrate a standard curve of glucose using the same assay was performed.

### Synchronous saccharification and capsular EPS production by *Cellulomonas flavigena*

The seed medium used was SCB hydrolysate obtained from preceding section, as well as the inoculum of 10% (v/v). M9 medium used for fermentation were composed of Na_2_HPO_4_.2H_2_O 4.2 g/l, KH_2_PO_4_ 1.5 g/l, NaCl 0.25 g/l, NH_4_Cl 0.5 g/l, yeast extract 0.5 g/l, 1M CaCl_2_.2H_2_O 0.01 % (v/v), 1M MgSO_4_.7H_2_O 0.2 % (v/v), trace elements (0.2%) EDTA 5g/l, FeCl_3_.6H_2_O 0.83 g/l, ZnCl_2_ 84 mg/l, CoCl_2_.2H_2_O 10 mg/l, H_3_BO_3_ 10 mg/l MnCl_2_.4H_2_O 1.6 mg/l along with 40g/l of sugarcane bagasse.

Fermentation was carried out till 30 hours with samples withdrawn at every two hours interval for growth studies and SRS (soluble reducing sugar) production from saccharification of sugarcane bagasse. Growth studies were performed using Triphenyl tetrazolium chloride (TTC) reagent which results in the formation of pink coloured product formazan if metabolically active cells are present in a culture (Defez et al., 2017). Absorbance of formazan measured at 480 nm was taken as a direct measure of growth. SRS production and cellulase activities were measured using DNS assay (Roy Chowdhury et al., 2011). For studying the production profile of extracellular polymeric substance (EPS) samples were collected at every 6 hours interval and subjected to extraction and purification with alkaline extraction and acid precipitation (detailed protocol given in later section).

Reducing sugar composition were analysed by HPLC, Agilent HPLC-1260 were equipped with Agilent Hiplex-H column with refractive index detector (RID). Both the column temperature and RID temperature were maintained at 35° C, flow rate of the mobile phase was kept at 0.6 ml/min with mobile phase as Milli Q water. Standards were run in order to identify the reducing sugars present in SCB hydrolysate. SCB hydrolysates of 0-30 hours were analysed by HPLC, for characterizing the reducing sugar composition throughout SDSF.

#### Extraction, Partial Purification and Structural characterization of EPS

For extracting water insoluble bacterial cell surface bound EPS, first the fermentation broth, cells of *Cellulomonas flavigena*, was filtered with a Whatman filter paper of pore size 8 μm to remove the insoluble bagasse particles. The retentate bagasse particles were washed twice and filtered to obtain the bound cells. All the filtrates were then centrifuged at 8000 rpm for 15 mins. Collected biomass in obtained pellets were treated with 1N NaOH under stirring condition for 2 hours. Then the solution containing the insoluble biomass and soluble EPS was centrifuged at 9000 rcf for 20 mins. EPS obtained in pellet were solubilized in 0.1N NaOH and then subjected to dialysis for 36 hours in a dialysis membrane composed of MWCO of 12 KDa. After dialysis the samples were freeze dried in Christ Alpha 1-2 LD plus free dryer with chamber pressure of 0.03 mbar and -55^°^C temperature till complete dryness was achieved. The fine powders obtained after freeze drying is the crude EPS, of which carbohydrate and protein estimation were done. Total carbohydrate and total protein estimation of EPS before and after dialysis were done by phenol sulphuric assay (Dubois et al., 1956) and BCA method respectively. Further deproteinization of the dialysed EPS was done to remove loosely bound proteins to check if any covalently bound protein is present in the EPS. Deproteinization was done using Sevag’s reagent and was repeated a no. of times till no protein in form of white layer as the interphase was obtained. Deproteinized samples were further dialysed to remove impurities generated during the deproteinization process.

Since our major focus is in the coating of microbial spores with crude EPS, in future we do not intend to separate and purify the exopolysaccharide (EP) from extracellular polymeric substance (EPS) and thus we do not perform biomineralization experiments with the deproteinised EPS.

#### Monosaccharide analysis by HPLC

Crude EPS samples in 5 mg/ml were hydrolysed using 4N TFA at 90-100^°^C in a silicone oil bath within a reflux apparatus. Hydrolysis were carried on till yellow to light brown color was achieved in the reaction mixture. After hydrolysis the hydrolysates were neutralized by constantly adding absolute methanol in an IKA-RV8 rotary evaporator. Neutralized samples were then dissolved in Milli Q water and analysed for the presence of monosaccharides, amino acids by High pressure liquid chromatography (HPLC) and thin layer chromatography respectively.

For HPLC of hydrolysed EPS and standard monosaccharides, were carried out using 100 % MilliQ water as the mobile phase. The Agilent HPLC (model no.1260) were equipped with Agilent Hiplex-H column with refractive index detector (RID). Both the column temperature and RID temperature were maintained at 35° C, flow rate of the mobile phase was kept at 0.6 ml/min.

Amino acids present in the EPS samples were identified using thin layer chromatography using mobile phase butanol: acetic acid: water 12: 3: 5 with 0.2 % ninhydrin as the detection agent. 20 amino acids standards of Sigma Aldrich were run along with the various concentrations of hydrolysed EPS. The R_f_ (retention factor) values of standards were compared with that of sample to determine amino acid residues present in EPS.

#### Surface charge determination and Molecular weight determination

Crude EPS were subjected to ion exchange chromatography to determine the overall charge of the biopolymer. The CM Sepharose column was equilibrated with 3-4 column volumes of 0M TRIS NaCl buffer pH 7.3-7.5. At first 1 mg/ml of crude EPS in TRIS NaCl buffer pH 7.3-7.5 were loaded on CM Sepharose matrix. After loading sample, step wise elution was carried on with 0, 0.2, 0.4, 0.6, 0.8, and 1M of Tris NaCl buffer. It must be noted that the concentration of Tris was maintained the same throughout as 20 mM only the concentration of NaCl was varied. All the eluted fractions were analysed using phenol sulphuric assay (PSA) method (Dubois et al., 1956). Similarly, sample was loaded on anion exchanger DEAE Sepharose matrix and sample elution performed in the similar manner. Along with carbohydrates samples were also analysed for proteins. For protein estimation BCA method was followed using BCA protein Assay Kit by Pierce. After analytical IEC, crude EPS was purified using preparative IEC to further perform gel filtration chromatography with the purified EPS.

For preparative IEC two column volumes of eluent were collected after loading sample and the buffer. These were then dialyzed for 24 hours to remove the excess salt present in mobile phase after which it was freeze dried to obtain powders of purified EPS fractions. These fractions are then re-dissolved in 0.1 N NaOH to perform gel filtration chromatography.

For gel filtration chromatography Sepharose CL 6B matrix was used as the stationary phase 0.1 N NaOH as the mobile phase. Void volume marker blue dextran was used along with standards of dextran 500KDa, 90 KDa, 60KDa, and 30KDa for standard curve preparation (Bhaumik et al., 2020). The standard curve was plotted by taking the logarithm of molecular weight on y axis and Ve /Vo in the x axis where, Ve is elution volume of a molecule and Vo is the void volume travelled by blue dextran. Purified sample from IEC was loaded on to gel filtration matrix eluted fractions were collected and analysed by PSA method.

## Results and Discussion

### Comparative analysis of pre-treatment by physico-chemical and microbial method

Functional groups present in untreated sugarcane bagasse (fig. 1) corresponds to wavenumbers 836, 897,1043,1160,1253, 1331, 1316,1370,1442,1513,1609,1728,2891,3413 cm^-1^. Among these the fingerprint region 800 to 1800 cm^-1^ mostly corresponds to functional groups present in lignin (Wittner et al., 2023). These vibrational bands were seen to be diminishing in case of all the combinations of treatment. 897 corresponds to C-H stretch of cellulose, 1023 to C-O stretch of cellulose and hemicellulose, 1112 vibration due to aromatic skeleton of lignin, 1160 to C-O-C vibrations of cellulose and hemicellulose, 1253 C-O stretch of lignin, 1331 C1-O vibrations in syringyl derivatives of lignin and C-H stretch of cellulose, 1370 C-H deformation of cellulose and hemicellulose. 1442 C-H deformation of lignin, 1513 aromatic skeletal vibration of lignin, 1609 aromatic skeletal vibration and C=O stretch of lignin, 1728 unconjugated C=O stretch of xylan, 2891 C-H stretch lignin, cellulose, hemicellulose, 3413 to O-H stretch of cellulose and hemicellulose. In case of chemical treatment majority of the lignin peaks was not seen except for 1630 corresponding to O-H and conjugated C=O of carbonyl group in lignin. Apart from that band at 897 cm^-1^ (C-H stretch of cellulose), was seen in all chemically pretreated SCB, unlike in bacterially digested SCB of 96 hours, this particular peak was seen to be diminished indicating saccharification by bacterial cellulase. However, in the bacterial digested samples obtained after 24 and 72 hours, the vibrational bands corresponding to lignin from 1111 to 1750 cm^-1^, were found, but with a much lesser absorbance than untreated raw sugarcane bagasse. These lignin specific bands, were completely diminished or got removed in bacterial digested samples after 96 hours. The FTIR spectrum shows effective delignification in all the sono-assisted alkaline pre-treated sugarcane bagasse samples as well as in bacterial digested SCB. However, saccharification of cellulose and hemicellulose can only be observed in bacterially digested samples. Although, vibrational band at 1043 due to C-O stretch of cellulose, hemicellulose (Asghar et al., 2015) as seen to be diminished in almost all the chemical and bacterial treated SCB samples. The reason can be the effect of sonication after alkaline treatment of SCB. The ultrasonic waves may have disrupted the polysaccharides present in SCB. Thus, it can be said that bacterial treatment is at par with that of chemical pre-treatment of sugarcane bagasse, moreover bacterial treatment not only removes lignin from LCB, but also simultaneously saccharifies the cellulose and hemicellulose moieties of LCB.

**Fig 1.**
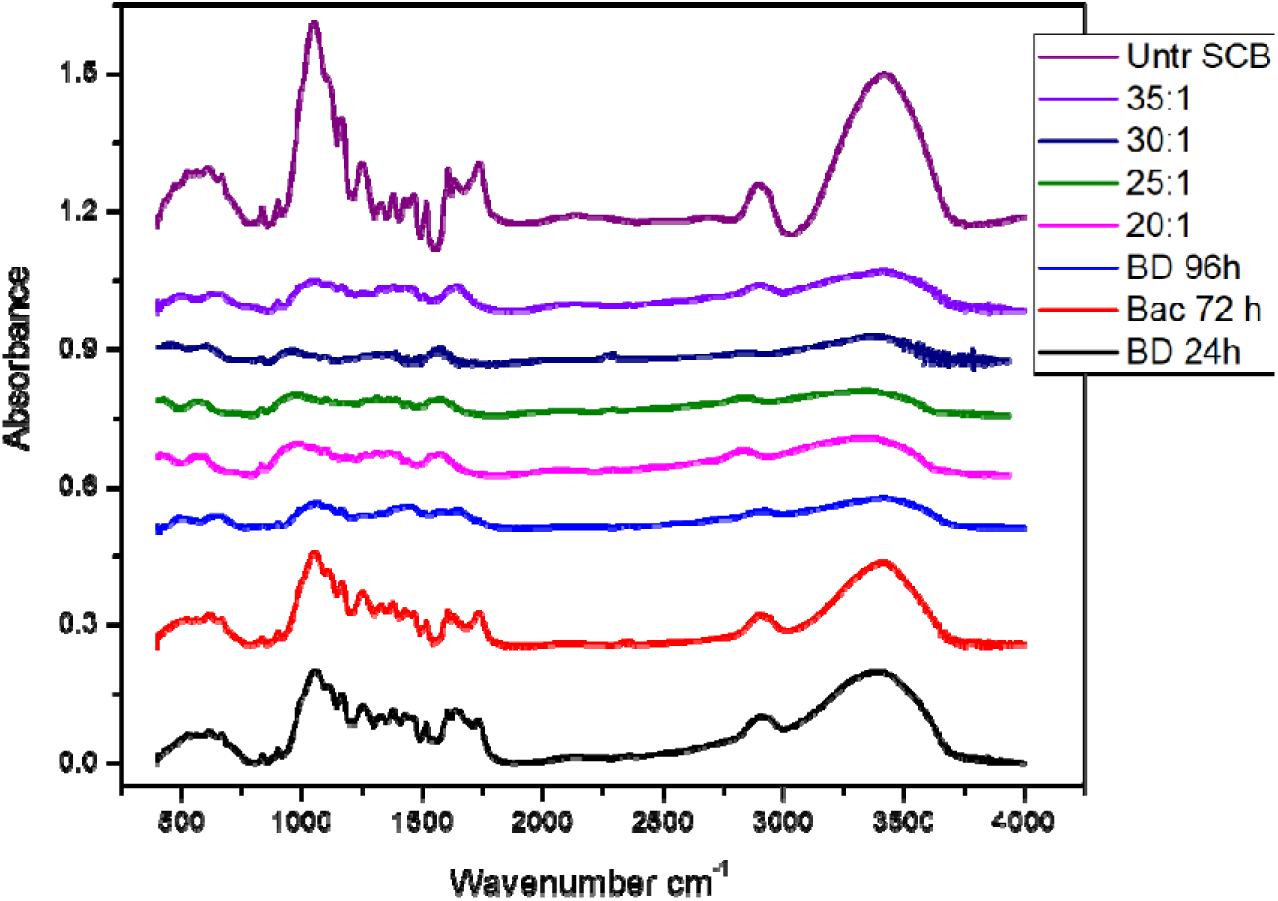
FTIR analysis of residue bagasse after sono-assisted alkaline pre-treatment and bacterial digestion method

The FTIR data (fig.1) can also be corroborated with the CHNS analysis (table 1), where it was seen that with both chemical and microbial treatment, the elemental composition of SCB changes. As observed from table 1, the carbon and nitrogen content were highest in the raw untreated SCB, which kept on decreasing with treatment. The nitrogen content got completely diminished in LSR 35:1 and 25:1 and in bacterial digested 96 hours. This indicates effective delignification in these combinations. Moreover, in the bacterial digested sample obtained after 96 hours, the carbon content also decreased from 46.4% to 22.6%, indicating the saccharification of the polysaccharides in SCB. While in bacterial digested sample obtained after 24 hours the carbon content decreased to only 43%. The carbon content in other chemically treated samples, LSR 20:1, 30:1 reduced very slightly as compared to bacterial treatment. Reduction in both carbon and nitrogen content in case of bacterial treated SCB indicates simultaneous delignification and saccharification of the biomass, unlike in chemical treatment where only the nitrogen content reduced due to delignification.

**Table 1.** CHNS analysis of Sono assisted alkaline pretreated, bacterial digested and untreated SCB.

| **Sample** | **C%** | **H%** | **N%** | **S%** | **O% +**<br>**inorganic**<br>**content** |
| --- | --- | --- | --- | --- | --- |
| **Raw**<br>**Untreated**<br>**SCB** | **46.381** | **7.097** | **4.321** | - | **42.201** |
| **LSR 33:1** | **35.57** | **6.431** | **0.00** | - | **58** |
| **LSR 30:1** | **46.501** | **8.141** | **1.456** | - | **43.902** |
| **LSR 20:1** | **45.935** |  | **0.639** | - | **53.426** |
| **LSR 25:1** | **40.97** | **6.47** | **0.00** | - | **52.56** |
| **Bacterial**<br>**digest 24 hrs** | **43.933** | **6.314** | **2.361** | - | **47.393** |
| **Bacterial**<br>**digest 96 hrs** | **22.6** | **8.394** | **0.24** | - | **68.766** |

X-ray diffractograms of SCB pre-treated by chemical and bacterial methods were analysed, (fig. 2a, b) to understand the change in crystallinity due to treatment. The major peaks of cellulose, at 002 plane crystalline cellulose 200 and 110 planes of amorphous cellulose were mostly prominent. The crystallinity of untreated SCB, was 38 % while it increased in case of all the pre-treated SCB samples. The sono-assisted alkaline pre-treated samples (fig. 2a), showed much higher crystallinity such as, in LSR 20:1 was 78%, in LSR 30:1 was 73%, in LSR 35:1 was 48%, the highest crystallinity in case of chemical treatment was observed in case 25:1, 97%. The bacterial digested (fig. 2b) or treated samples had crystallinity, in the range of 68% for 24 hours digested samples, 76% for 72 hours, 92% for 96 hours digested samples. It is worth mentioning that the bacterial digestion or treatment of SCB was well comparable with the chemical treatment. This is because the increase in crystallinity of bagasse was highest in LSR 25:1 and bacterial digested 96 hours samples. This hike in crystallinity indicates effective delignification as well removal of hemicellulose, which contributes to amorphous fraction of LCB. The elimination of lignin and hemicellulose exposes the crystalline cellulose, that causes the increased crystallinity with pre-treatment (da Costa et al., 2023).

**Fig 2.**
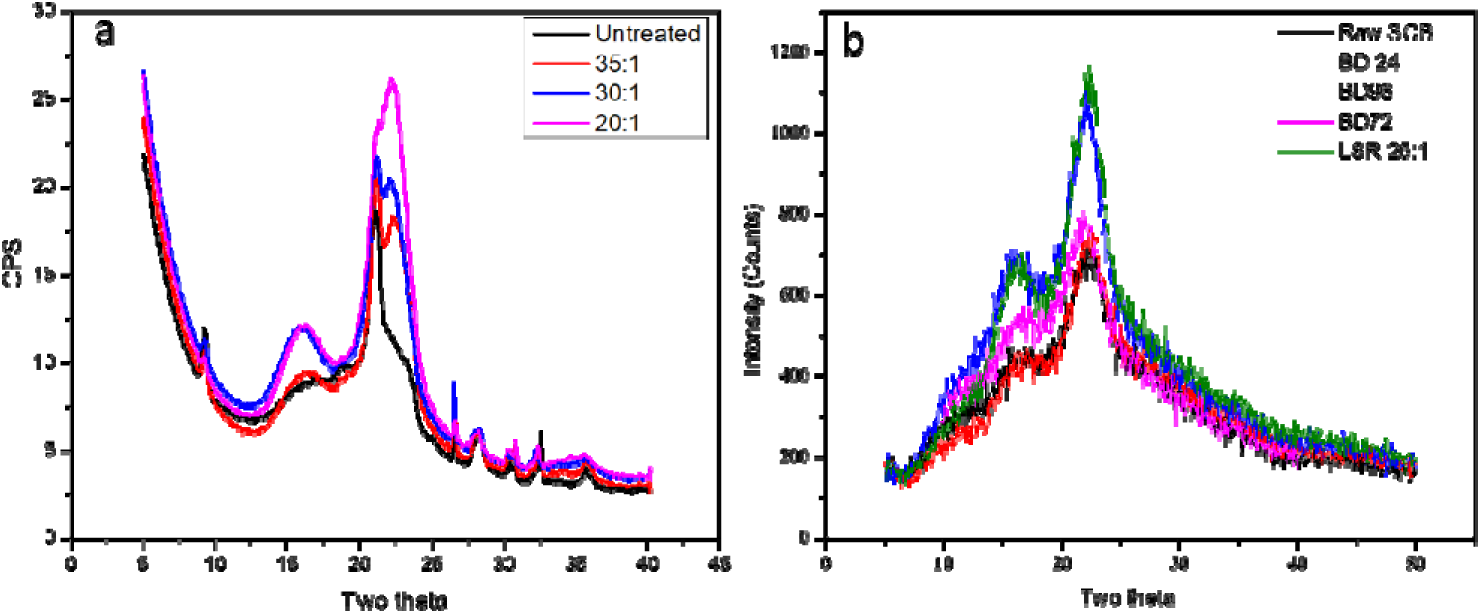
XRD analysis of residue bagasse (a) after sono-assisted alkaline pre-treatment (b) bacterial digestion method

### Morphological characterization of chemically pretreated, untreated and bacterial digested sugarcane bagasse

The effect of pre-treatment both chemical and biological were also evaluated morphologically through scanning electron microscopy. As observed from the SEM images (fig 3a-c) of the transverse sections of sono-assisted alkaline pre-treated SCB versus control untreated SCB, disruptions in the cell wall structure of LCB is quite prevalent. In the control untreated SCB (fig. 3c), the intact vascular bundle and fibre lumens can be seen, which got largely degraded in the physio-chemically treated samples of SCB. Pores in the vascular bundles can be well observed in the treated samples (fig 3a, b), indicating the removal of lignin and hemicellulose moieties from SCB. This can also be corroborated with the XRD data (fig. 2b), where LSR 25:1 showed very high CrI, majorly due to the deletion of the amorphous lignin and hemicellulose from cell wall of SCB. Moreover, the bacterial digested (BD) SCB samples (fig 4a-j) showed extensive disruptions of the vascular bundles as opposed to bundles of raw untreated bagasse. In addition to that, with increase in amount of bacterial digestion time, the disruptions of the cell wall structure of SCB largely increased. With higher magnification, in control as well as BD samples, the effect of pre-treatment can be better seen. As can be observed, in BD 24 (fig 4b, g), pores or cavities were formed after treatment, BD 72 (fig 4c, h) showed much larger pore or roughness of the cell wall, BD 96 (fig.4d, e, j) showed complete unevenness or damages in the intact vascular bundles present in cell wall of SCB.

**Fig 3:**
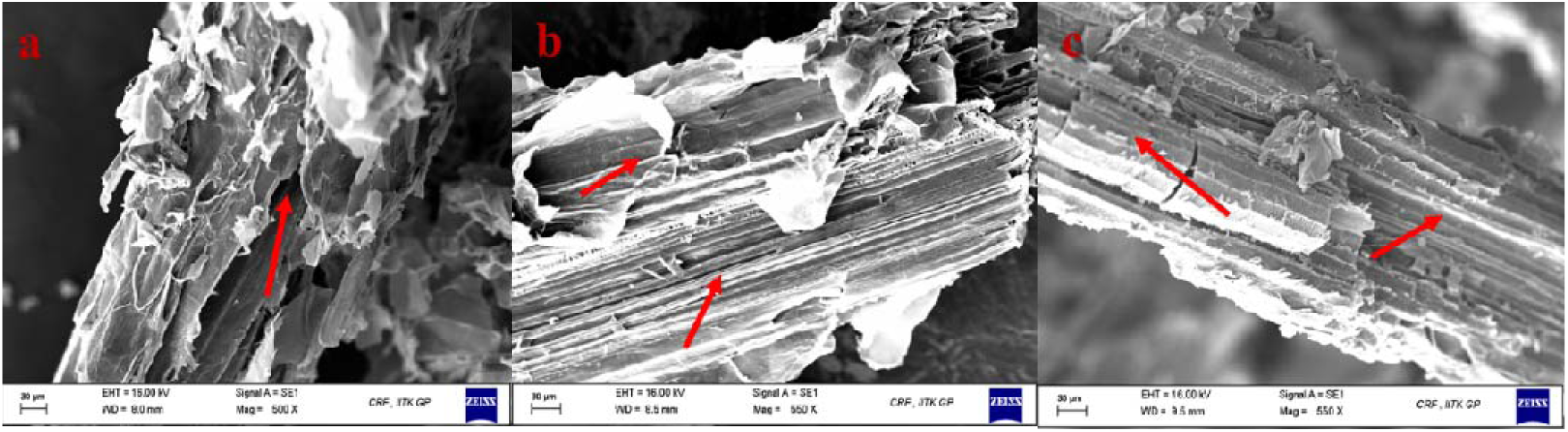
(a, b) sono assisted alkaline pretreated sugarcane bagasse of LSR 25:1 (c) raw untreated sugarcane bagasse; Red arrows indicating structural changes of sugarcane bagasse after physico-chemical pre-

**Fig 4:**
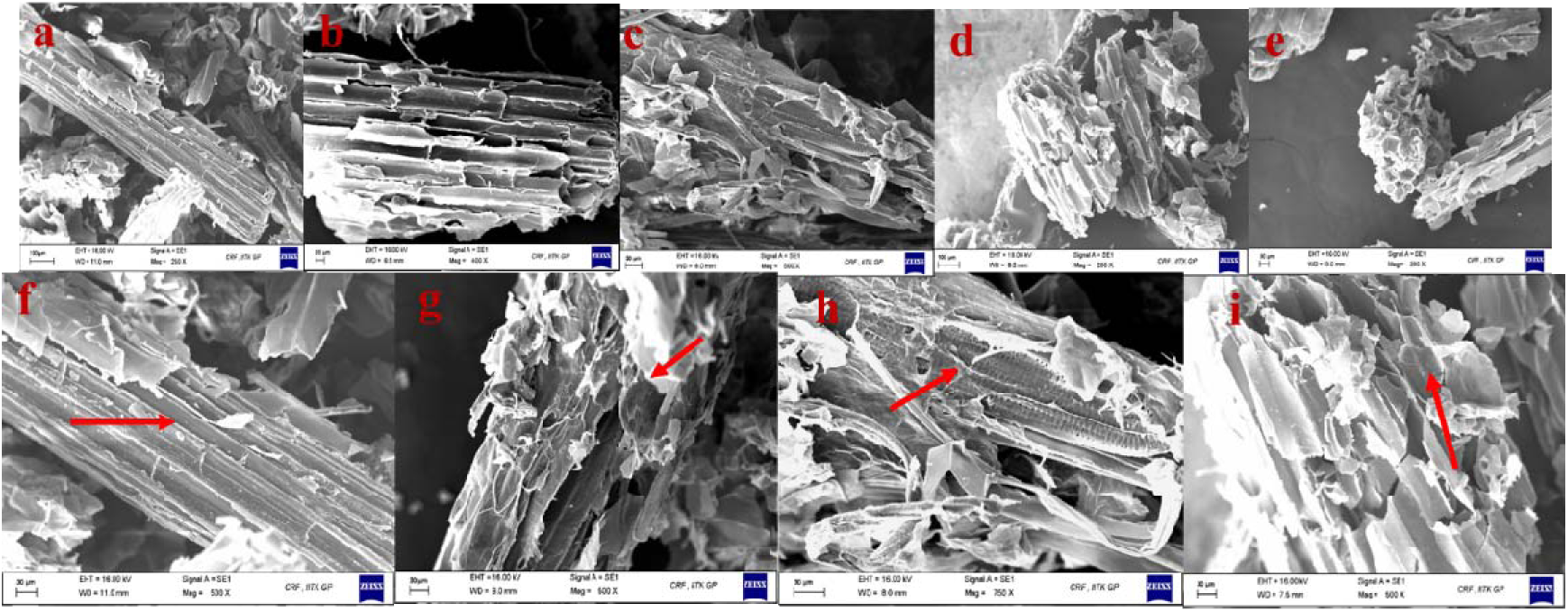
(a, f) untreated SCB, (b, g) BD 24 hours, (c, h) BD 72 hours (d, c, j) BD 96 hours

Morphological characterization of pre-treated sugarcane bagasse is essential to prove the disruption or damage of cell wall due to treatment with removal of lignin and hemicellulose. The SEM images shown in this study, for both the type of pre-treatments, comply much with literature, where disturbances in the intact vascular bundles are reported as a proof of effective delignification and hemicellulose elimination (Thite and Nerurkar, 2019; Yahyaee et al., 2022).

### Microbial growth on pretreated and untreated sugarcane bagasse

The growth of *Cellulomonas flavigena* (CF) cells using both chemically pretreated as well as untreated sugarcane bagasse were studied to understand whether pre-treatment is at all required by the CF cells to perform saccharification of the cellulose present in SCB. Here untreated sugarcane bagasse is basically the bagasse subjected to SDF or biological treatment by the growth of CF cells. produced by growth of *Cellulomonas flavigena* on untreated and pretreated SCB. From fig 5a, similar growth profiles of CF cells in both chemically treated and untreated bagasse, indicating that CF cells are competent in growing raw SCB. In fact, it was seen that the cells are growing at a much faster rate with greater cell density (since OD@480nm corresponds to the metabolically active cells) on the untreated SCB. Thus, it can be said that the SCB can be directly used by the CF cells for the production of value-added products without the hassle of cost incurred in expensive chemical pre-treatments followed by enzymatic saccharification.

**Fig 5:**
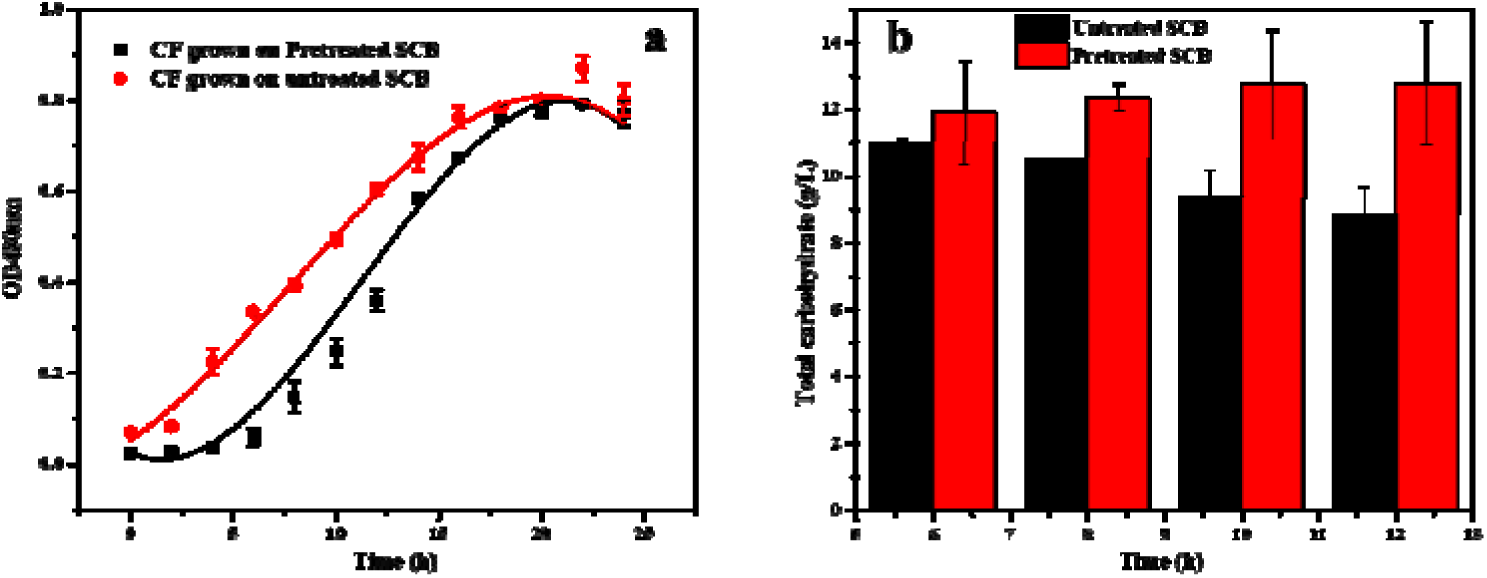
(a) Growth of *Cellulomonas flavigena* on pretreated and untreated SCB (b) Total carbohydrate

However, the on monitoring the total carbohydrate concentrations of culture supernatants from both pre-treated and untreated biomass, it was found that higher amount of carbohydrates were generated by the CF cells, when grown on sono-assisted chemically treated SCB of LSR 25:1. The highest total carbohydrate production (fig. 5b) was 12.77 ± 1.8 g/L, after 12 hours of fermentation, in case of pre-treated biomass, while that in case of untreated one was 11 ± 0.15 g/L after 6 hours of fermentation. It can be seen from fig 5b that the total carbohydrate content decreased with fermentation time in case of untreated bagasse. One possible reason can be the larger accessibility of cellulose to enzymatic digestion in pre-treated bagasse, rather than in untreated biomass. Moreover, since higher amount of cellulose is available for cell growth, so the cell utilizes the polysaccharides to produce reducing sugars, whereas in case of untreated bagasse, lesser amount of cellulose is accessible so the cell utilizes smaller amount of cellulose for saccharification and further thrives on the reducing sugars produced out of SDSF. As observed from fig.6a, reducing sugar produced by the CF cells shows an increasing profile with time. This indicates that as the SDSF time increases, the CF cells grows actively by utilizing cellulose crystals to produce reducing sugars like cellobiose, glucose and xylose (fig 6b). The reducing sugars obtained at 12^th^ hour, were identified after comparing the retention times with that of standard mix, containing cellobiose, glucose and xylose (CGX). From HPLC results the cumulative concentration of reducing sugars was obtained as 9.6 g/L, differing slightly from DNS results of 8.8 g/L. Since DNS is a colorimetric assay, which is much more prone to erroneous results as compared to HPLC producing very high precision results, so the reducing sugar concentration of 9.6 g/L is considered thereafter.

**Fig 6.**
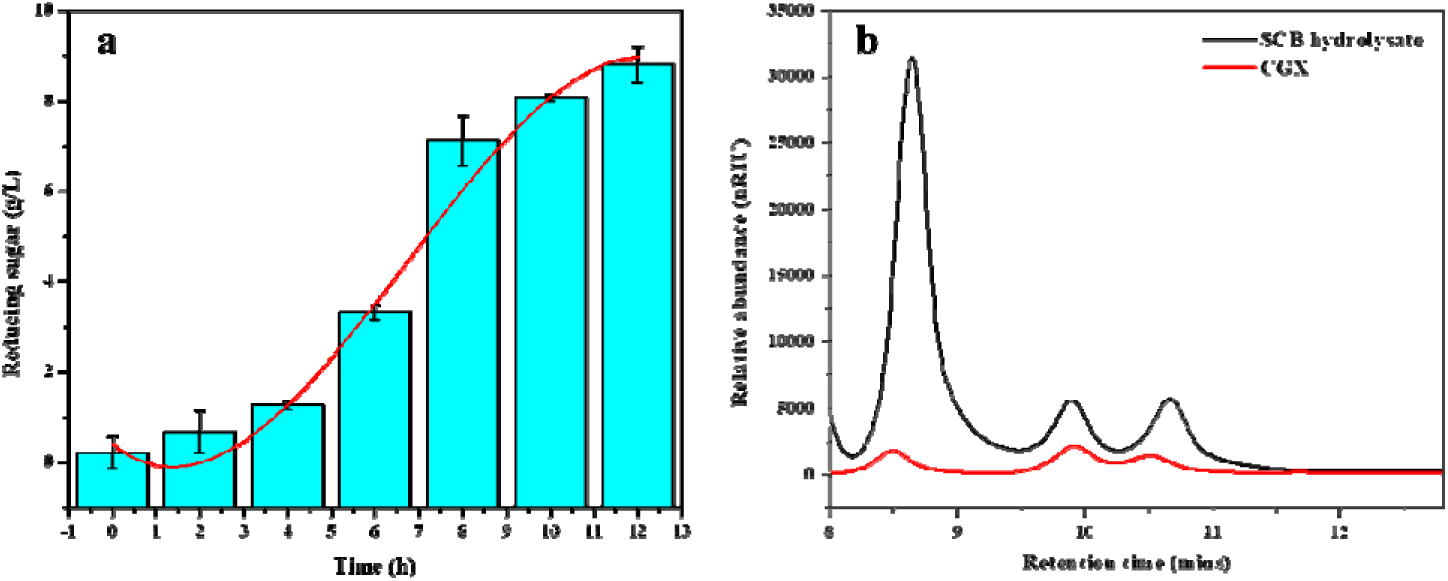
(a) Reducing sugar content produced by CF by growing of untreated SCB (b) HPLC chromatograms showing the composition of reducing sugars present in the SCB hydrolysate along with standard mix containing reducing sugars cellobiose, glucose and xylose in order.

#### Simultaneous saccharification and fermentation for production of EPS

Fermentation of waste sugarcane bagasse by cellulolytic bacterium *Cellulomonas flavigena* resulted in the production of 4.04 g/l of extracellular polymeric substance (fig. 7a). As observed from fig.7a the cells start to grow exponentially from 6 hours of fermentation however it can be observed that the production of reducing sugar starts within the early log phase of growth. The log phase can be seen from 6 to 22 hours (fig. 7c) after this phase, it can be observed that the reducing sugars produced by the saccharification of sugarcane bagasse were partially utilized by the cells. It can be observed that, from 12-16 hours there is not much production of reducing sugars, which indicates the consumption of the same by the cell. There is a constant saccharification of cellulose and hemicellulose present in SCB along with constant utilization of reducing sugars during the log phase. The production of reducing sugar comes to a halt during mid log phase as the cells are actively feeding upon the sugars that are formed out of saccharification, as a result of high cellulase activity during this phase (fig. 7a). As the cells gradually starts moving towards late log phase there is again a rise in the cellulase activity with increasing yield of reducing sugar. As a result of which a steep increase in cell growth can be seen from 24 to 26^th^ hour of the growth curve can be observed. After 26^th^ hour a fall in the growth curve can be seen along with decrease in the concentration of reducing sugar and cellulase activity (fig. 7a, c). The EPS production can be seen to start from 10^th^ hour with maximum production at 18^th^ hour. There is a clear indication that, EPS is a growth associated product as it mostly accumulates during the log phase and there is a sharp decline after that. fig. 7a shows HPLC chromatograms of all the time points from 0-30^th^ hour showed reducing sugar composition of culture supernatant (SCB hydrolysate) as cellobiose, glucose and xylose, when compared with standard mix of these three reducing sugars. Constant saccharification and fermentation can be also be well corroborated with the HPLC chromatograms of various time intervals (fig. 8a). Appearance of cellobiose followed by the occurrence of glucose and xylose is an indication of hydrolysis of complex substrates like cellulose and hemicellulose (fig. 8b).

**Fig 7.**
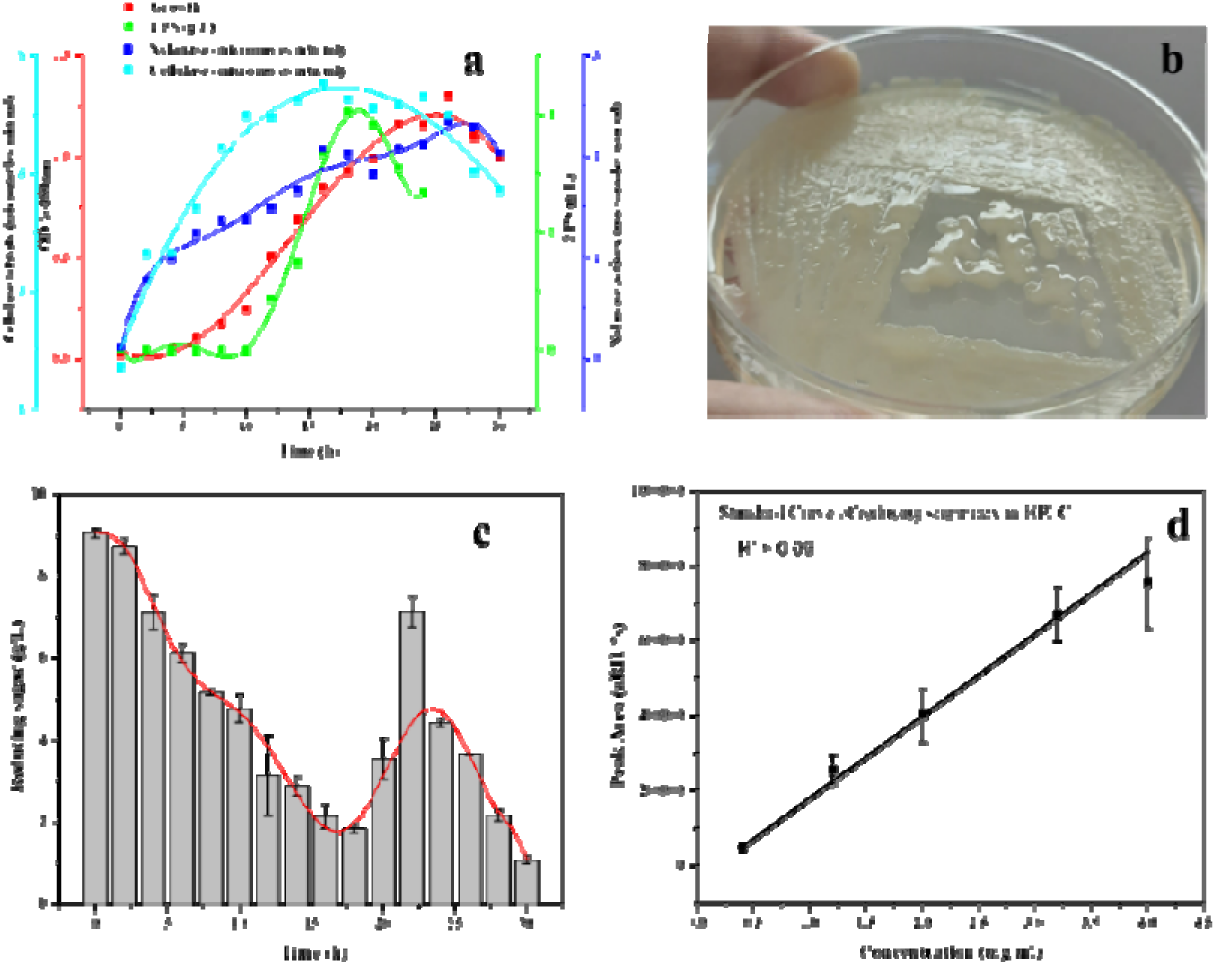
(a) Time course growth, EPS production profile and Cellulase activity of *Cellulomonas flavigena* grown on sugarcane bagasse (b) Colonies of *Cellulomonas flavigena* producing slime like EPS extracellularly (c) Time course reducing sugar profile of *Cellulomonas flavigena* grown on sugarcane bagasse (d) Standard curve of reducing sugar mix in HPLC All the experiments were performed in biological replicates of three (*n*=3)

**Fig 8.**
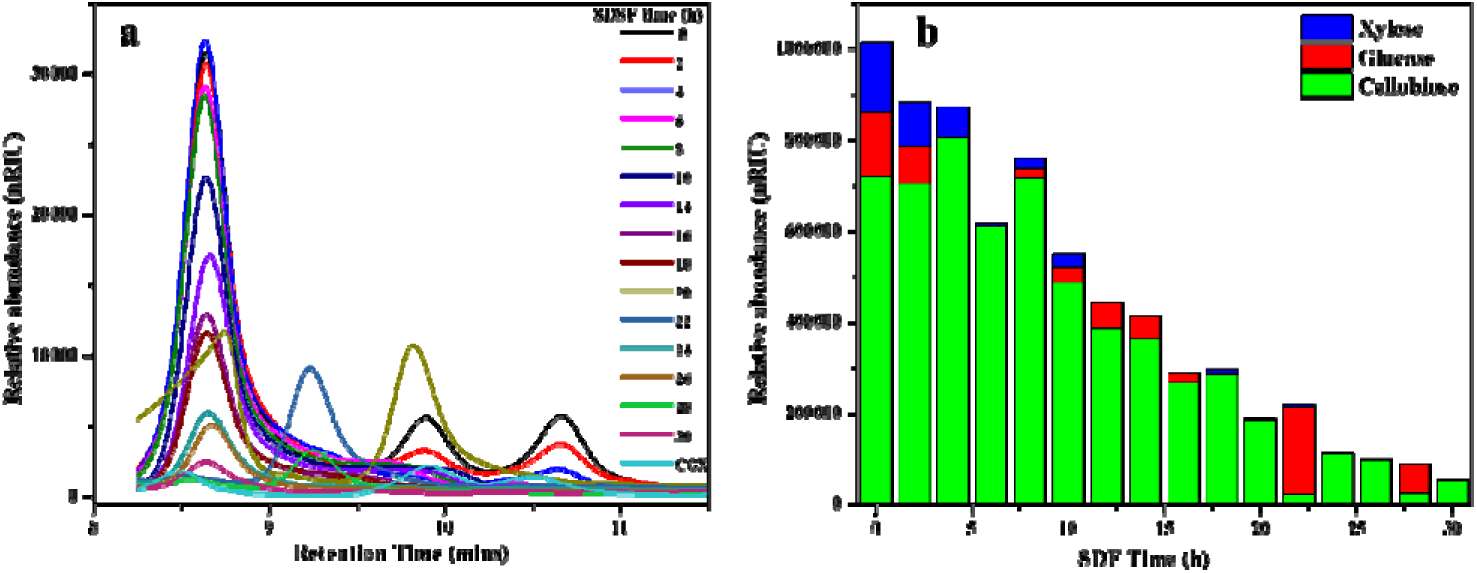
(a) HPLC chromatograms of SCB hydrolysates at different time intervals of simultaneous and fermentation of sugarcane bagasse (b) Differential proportion of each reducing sugar in SCB hydrolysate with time.

In this study, an interesting profile of the soluble reducing sugar were obtained during 30-hour long saccharification and fermentation. As observed, log phase can be captured when the cells are directly fermenting on the reducing sugars produced during the lag to log phase transition. This means that the CF cells accustoms itself in the growth media conditions by expressing cellulase and xylanases for hydrolysing cellulose and hemicellulose in order to grow and survive with the available nutrients. On saccharification of these polysaccharides the bacterium feeds on to the monosaccharides and disaccharides like glucose, xylose and cellobiose (fig. 8b) respectively. With time, as the cells grow they utilize these reducing sugars for the synthesis of different primary and secondary metabolites. EPS mostly falls under the category of secondary metabolites which is not directly required by for the cell’s survival. So, these biological macromolecules at the cell’s exterior starts to accumulate, only when a nutrient deprivation like condition is encountered by the cells, they breakdown these complex polymeric molecules into simple monomers for its survival (Costa et al., 2018). This strategy of survival by CF cells can be also be seen this study (fig. 7a), after maximum accumulation of product when stationary phase is achieved these macromolecules are drastically utilized by the cells. Also, during this phase, a hike in reducing sugar concentration can be observed, which again can be a survival strategy of CF cells to saccharify the residual number of polysaccharides present in sugarcane bagasse. Cells were found to hydrolyse or utilize both cellulose and hemicellulose as reducing sugars like cellobiose, glucose and xylose were obtained in the culture supernatants when maximum saccharification occurred. Proportions of each reducing sugar also varied with the progression of SDSF (fig 8b), which is indication of the cell’s preference of carbon source for their survival. It was indeed interesting to observe and can be further analysed to understand the underlaying mechanism of optimal growth of CF cells towards particular reducing sugar.

Carbohydrate and protein fractions of EPS were studied before dialysis, after dialysis and after deproteinization. It can be observed from table 2 that carbohydrate fraction increases in each step, some of the loosely bound proteins were still present after dialysis which were removed by repeated deproteinization. Even after performing deproteinization multiple times, it can be observed that proteins are present in the EPS. The quantity of protein content in EPS reduced to almost half after several deproteinization steps (Table 1). This indicates that these proteins are covalently bound which can be not be separated by organic solvent treatment present in Sevag’s reagent.

**Table 2:**
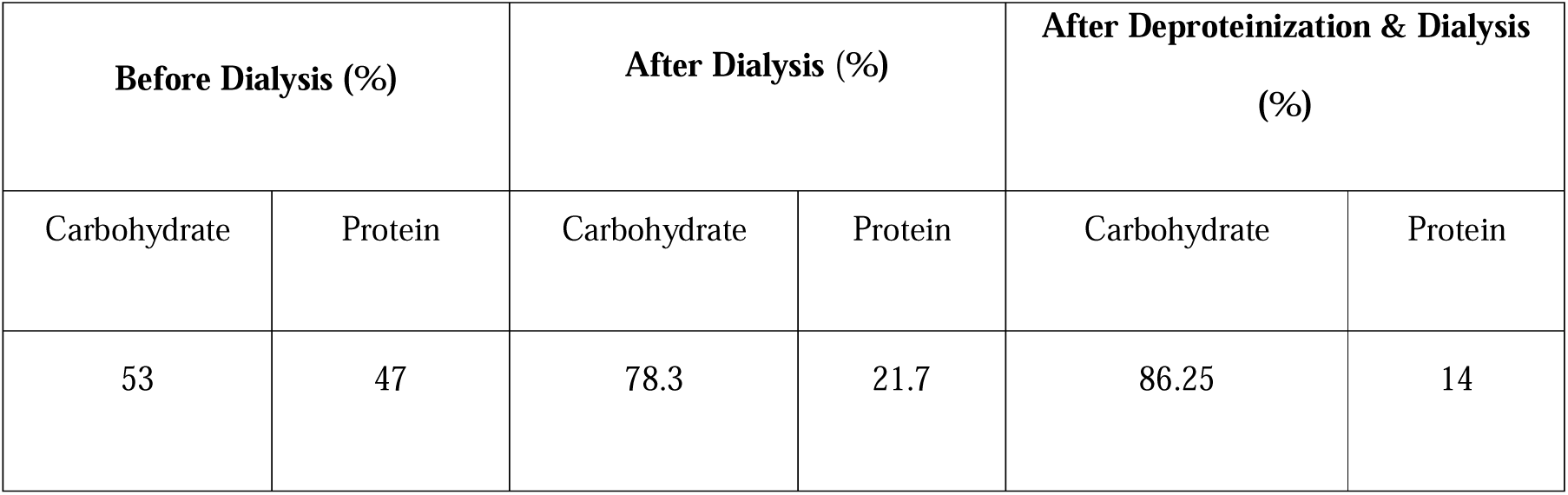
Carbohydrate and protein quantification in EPS.

| Before Dialysis (%) |  | After Dialysis (%) |  | After Deproteinization & Dialysis (%) |  |
| --- | --- | --- | --- | --- | --- |
| Carbohydrate | Protein | Carbohydrate | Protein | Carbohydrate | Protein |
| 53 | 47 | 78.3 | 21.7 | 86.25 | 14 |

**Table 3:** HPLC results of crude EPS hydrolysate and standard monosaccharides.

| Retention time | Peak area % | Monosaccharide |
| --- | --- | --- |
| 6.9 | 3.78308e4 | D- Mannosamine hydrochloride |
| 10.535 | 8.08245e4 | D -(+) Xylose |
| 7.013 | 3.13250e5 | Crude EPS |
| 10.545 | 6.39034e5 | Crude EPS |

**Table 4:** TLC results of crude EPS hydrolysate and standard amino acids.

| R<sub>f</sub> value | Amino acids |
| --- | --- |
| 0.34 | L-Glutamine |
| 0.61 | L-Tyrosine |
| 0.35, 0.62 | EPS hydrolysate |

### Monosaccharide analysis: HPLC / Monomeric composition of EPS

The chromatogram obtained from HPLC (fig. 9b) showed the presence of two peaks well separated from each other. After comparing the retention times of various monosaccharides with the hydrolysed EPS it can be observed that peak at 7.00 minutes corresponds to the amine sugar D-Mannosamine hydrochloride and at 10.53 corresponded to D (+) Xylose. The Exopolysaccharide part of EPS was found to be heteropolysaccharide comprised hexose amine and pentose sugars.

**Fig 9.**
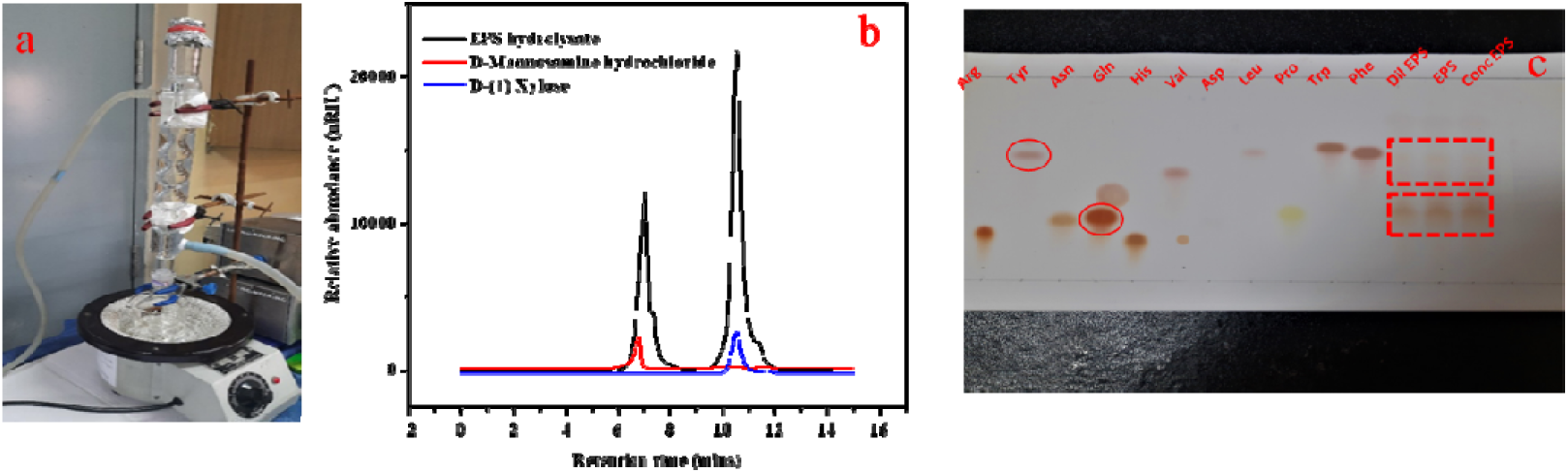
(a) Acid hydrolysis of crude EPS using reflux apparatus (b) HPLC chromatogram showing monosaccharide composition of EPS (c) TLC of crude EPS detecting amino acids in the EPS hydrolysate

Xylose and mannose present in the EPS can be due to the utilization of hemicellulose of sugarcane bagasse by *Cellulomonas flavigena*. These mannose residues obtained from metabolizing hemicellulose possibly derivatized by cells to form amine sugars, mannosamine, which is a component of glycoproteins present in cell surface S layers (Schäffer and Messner, 2004). Cell surface glycoproteins present in bacteria and yeast are known to be composed of rare sugars like mannosylated glycans or N-acetyl monoamine, N-acetyl glucosamine (Hirayama et al., 2019; Schäffer and Messner, 2004). Moreover, Schäffer and Messner, (2004) in their study showed the presence of O-methyl group capping on the surface glycoproteins in bacteria. FTIR spectrum shows the presence of methyl group (data not shown; submitted elsewhere). These further indicates that target EPS in this study has a glycoprotein like nature as can be confirmed by ion exchange chromatography, HPLC, TLC data (fig. 9, 10).

Along with the monosaccharide composition of EPS amino acid composition were analysed using Thin layer chromatography (fig 9c). After comparing the R_f_ values of standards with sample it can be found that the amino acid residues present in EPS are L-Glutamine and L-Tyrosine. Glutamine and tyrosine are both polar neutral molecule present in the EPS.

Presence of amine sugars and polar neutral amino acid residues in EPS might indicate the dominance of positively charged and neutral fractions of EPS obtained from IEC. Since the main target of this study is to form a bioconjugate of EPS with bacterial spores for application in biomineralization, EPS in its crude form has only been characterized. The crude EPS is essentially a mixture of polymeric substances like polysaccharides, polypeptides, on hydrolysis of the crude EPS the monomeric components obtained are mostly monosaccharides and amino acids. The monomeric composition can be corroborated with the structural insights obtained from FTIR spectra. Presence of amide bonds in FTIR spectra (data submitted elsewhere) can be justified with the presence of polypeptide like components in the EPS. Amine group can be also be found from FTIR data which is most likely due to the presence of mannosamine sugars in the polysaccharide moiety of EPS, which can be confirmed by HPLC. Also, presence of aldehyde groups and large no of OH groups in EPS can be corroborated with presence of pentose and hexose sugars in the polysaccharide chain of EPS as observed from HPLC. Presence of β glycosidic linkage was seen from the FTIR spectra (data not shown; submitted elsewhere), which possibly can be the linkage between mannosamine and xylose residues forming the polysaccharide backbone of the EPS.

### Purification, surface charge and molecular weight determination

Crude EPS was subjected to both purification and analysis by ion exchange chromatography. Since crude EPS consisted of both carbohydrates and proteins so by performing ion exchange chromatograhy both the biomacromolecule fractions can be identified.

As observed from fig. 10 a, the major fraction of the crude EPS eluted when 0.4 M NaCl Tris was passed through the CM Sepharose column. Minor peak at 0.8 M fraction can be also be seen. Another major fraction can be also be seen being eluted with 1 M concentration of NaCl Tris buffer. Since a major peak was obtained at 0.4 M fraction of NaCl Tris, this fraction was considered to contain the maximum EPS content and so were further studied in gel filtration chromatography. Fig 10b shows the elution of proteins with 0.8 M NaCl Tris in CM Sepharose matrix. Mostly one major fraction of protein can be seen present in the EPS, exhibiting strong positive charge that eluted with a relatively high concentration of NaCl Tris buffer. Also, at 0.8 M Tris NaCl concentration it can be seen that both proteins and carbohydrates are eluting out with protein having a major peak and carbohydrate with a minor peak. This suggests that the EPS is a glycoprotein like molecule which can be proved further in this study. Interestingly in DEAE sepharose matrix fig. 10c as well few carbohydrate fractions of EPS can be seen to be eluted although a major fraction was obtained with 0 M NaCl-Tris buffer. This corresponds to the unbound molecules to DEAE matrix, which mostly comprises of neutral and positively charged molecules. Two fractions of EPS can be seen to be eluted with 0.8 M and 1 M NaCl Tris in DEAE Sepharose matrix. This indicates that the EPS is mostly composed of positively charged residues with some fractions exhibiting negative charge as well. Fig. 10d represents anion exchange chromatography of EPS with a stepwise elution using a concentration gradient of NaCl, where anionic proteins components have been analysed. As can be observed two peaks have been detected one corresponding to neutral or cationic proteins and the other anionic proteins. From fig. 10 b it can be seen that the major protein component is cationic in nature, which most likely has eluted at 0 M NaCl during anion exchange chromatography. Moreover, protein fraction eluted at 0.8 M NaCl concentration has a much lower concentration in anion exchanger than protein fraction eluted at the 0.8 M NaCl concentration of cation exchanger. This is indicative of the fact that the protein moiety present in EPS has more no. of positively charged residues than anionic residues.

**Fig 10:**
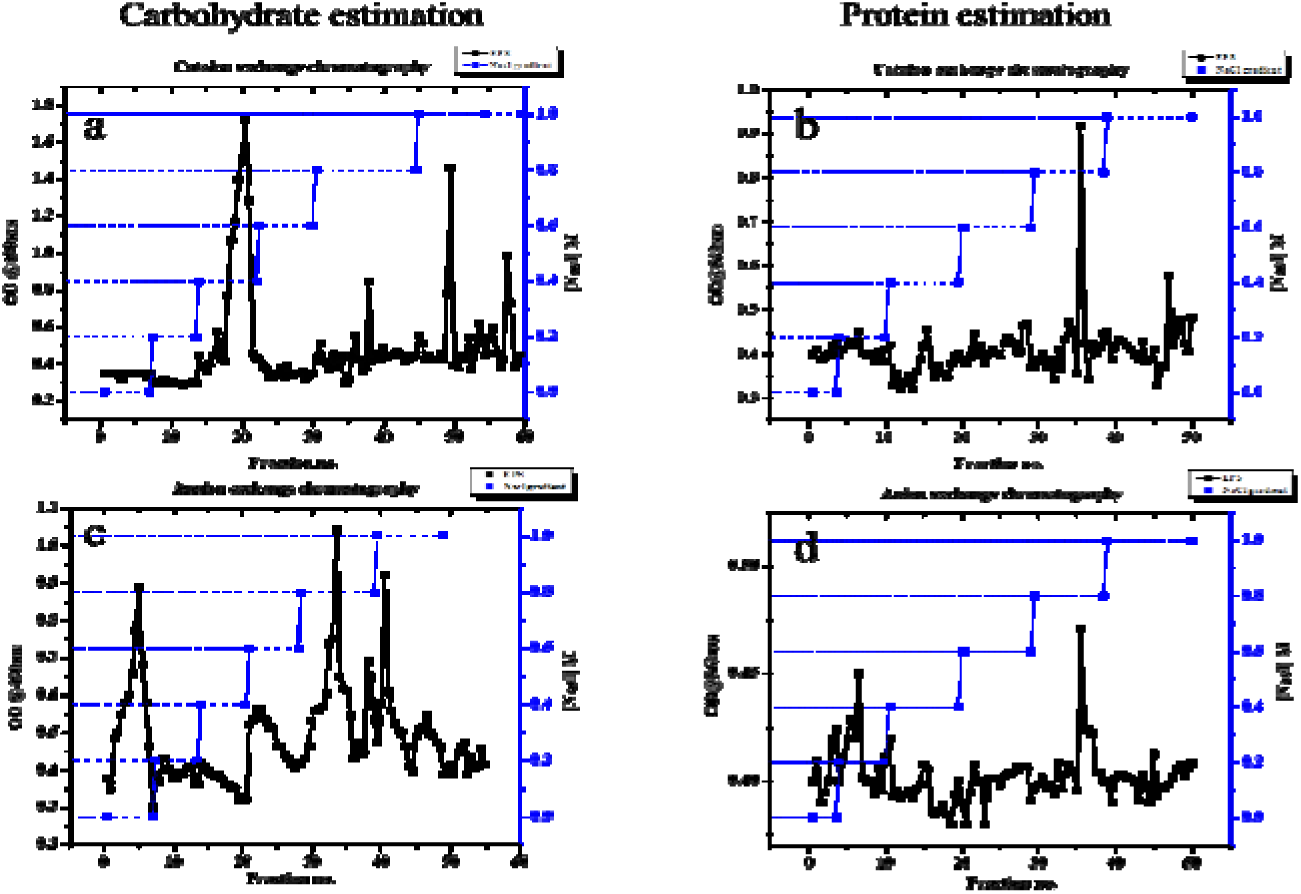
Stepwise elution of crude EPS with a NaCl Gradient 0 to 1M Tris NaCl buffer pH 7.3 (a). in CM sepharose matrix with carbohydrate estimation (b) in CM sepharose matrix with estimation protein (c) in DEAE sepharose matrix with carbohydrate estimation (d) in DEAE matrix with estimation protein

It must be noted that the EPS is a complex polymeric substance made up of a network polypeptides and polysaccharides which often bears ionic characteristics. The ion exchange chromatograms that were obtained in this study are manifestations of diverse chemical and ionic properties of bacterial extracellular polymeric substance. Purified fraction (0.4 M) from ion exchange chromatography were analysed to determine the molecular weight. The molecular weight of the EPS was obtained from the standard curve of dextran. As observed from fig. 11b there are two major peaks in the gel filtration chromatogram of EPS, one has high molecular weight of 237 KDa and the other 29 KDa. These fractions are obtained after further purification of the 0.4 M fraction from IEC.

**Fig 11.**
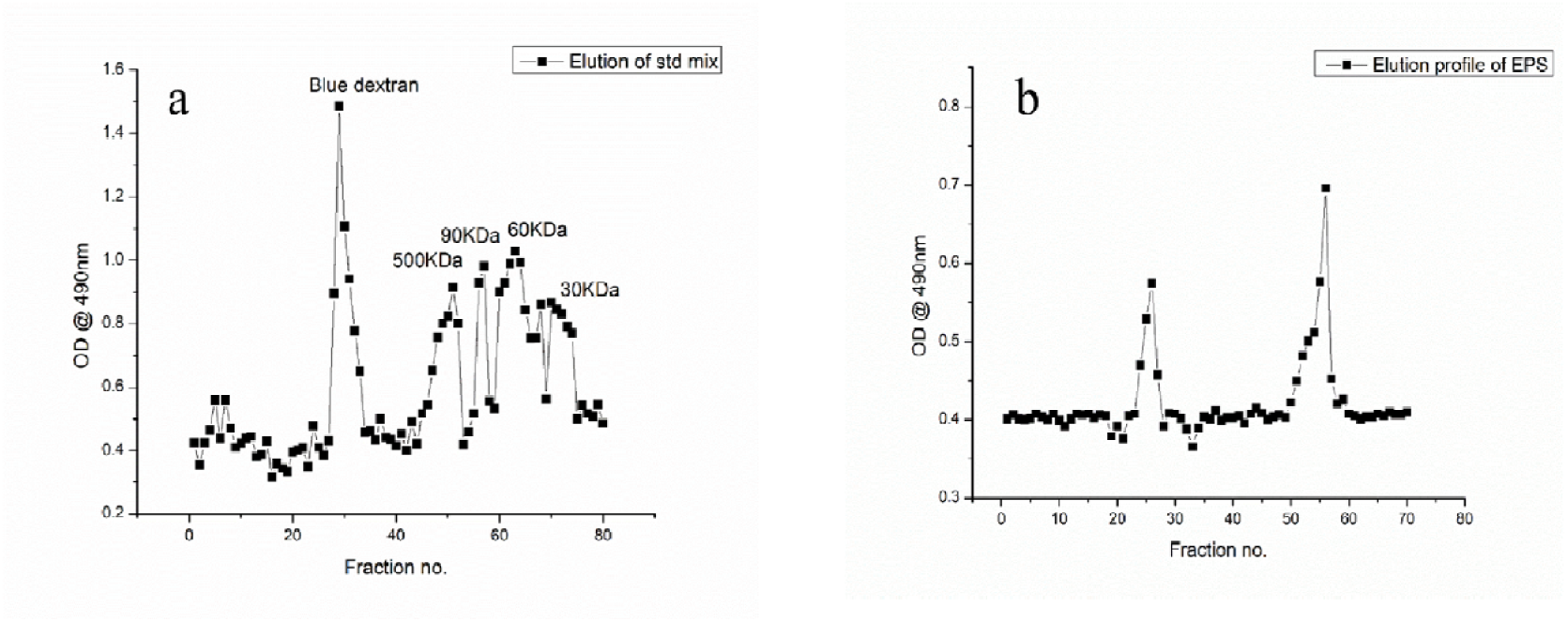
Gel filtration chromatograms of (a) standard mix showing elution profiles of blue dextran (void volume marker) then various molecular markers of dextran in descending order of their sizes (b) elution of profile of purified EPS from IEC showing two major fractions of the molecule

Most possible reason behind the occurrence of multiple fractions in IEC & GFC can be due to presence of various polymeric fractions within the EPS (Ajao et al., 2021; Dong et al., 2020; Yuan et al., 2020). These polymeric fractions might be of the same nature and composition but of diverse molecular sizes due to their different stages of synthesis. Nevertheless, these facts need to be proved further in order to have solid conclusion regarding the various polymeric structures of this EPS, which is beyond the scope of this study.

## Conclusion

Sugarcane bagasse is a major agro-industrial waste of countries like India, Brazil etc, and thus this waste is abundantly available. This waste is rich in cellulose, because of which it has been widely used for biofuel production. However, the pertaining problem of high cost involved in lignin removal from lignocellulosic biomass is a critical limitation for the utilization of this biomass in biotechnological applications. Literature extensively deals with chemical and biological pre-treatment in tandem for delignification and hemicellulose removal, in order to increase the accessibility of cellulose for saccharification thereby yielding reducing sugars for fermentation for biofuel production. In our study, we have proposed a unique simultaneous delignification saccharification and fermentation (SDSF) strategy by *Cellulomonas flavigena* cells. These cells are utilizing the cellulose crystals of raw sugarcane bagasse by simultaneously removing the lignin and hemicellulose moieties from the LCB. The cells during their growth in SCB, produces a hydrophobic biopolymer (EPS), which is a value -added product that can be further applied in various environmental and biomedical applications. A comparative analysis of sono-assisted vs microbial pre-treatment shows effective delignification and hemicellulose removal in LSR 25:1 and bacterial digestion of 96 hours. Apart from that, these bacterial cells produced a water insoluble biopolymer, composed of monosaccharides like xylose, mannosamine and amino acids like L-Glutamine, L-Tyrosine. The biopolymer had a glycoprotein like nature with positively charged molecular weight fractions of 29KDa and 237KDa. The waste valorised biopolymer produced by *Cellulomonas flavigena* would be further utilized for coating bacterial spores for application in self-healing of concrete.

## Author’s contribution

AD: experimentation, analysis, writing original draft; SB: Formal analysis, experimentation, writing; RK: Supervision, reviewing draft

## Acknowledgement

Central Research Facility and Department of Bioscience and Biotechnology, IIT Kharagpur are duly acknowledged for all the instrumentation facilities. AD acknowledges DBT, Govt. of India for financial assistance, SB acknowledges MNRE, Govt. of India for financial assistance.

